# Conformational analysis of liganded human hemoglobin by cryo electron microscopy

**DOI:** 10.1101/2025.07.07.661630

**Authors:** Katsuya Takahashi, Yongchan Lee, Tomohiro Nishizawa, Jeremy R. H. Tame

**Affiliations:** Graduate School of Medical Life Science, Yokohama City University, Suehiro 1-7-29, Yokohama, Japan

**Keywords:** allostery, oxygen binding, evolution, cryo-electron microscopy, crocodile

## Abstract

The long-standing debate on the preferred conformation of liganded hemoglobin (Hb) in solution has yet to be completely resolved. While some studies have used lyophilized human hemoglobin for structural studies by cryo-EM, we recently presented the first cryo-EM analysis of freshly prepared human and crocodilian Hbs. Further three-dimensional (3D) classification analysis of these datasets reveals distinct structural characteristics. CO-bound adult human Hb (CO-HbA) shows a mixture of conformations, with the R2 conformation most populated, R strongly represented, and other intermediate states present in sufficient quantity to produce maps. CO-bound crocodile Hb showed the R conformation and, unexpectedly, a smaller population of molecules in a T-like conformation. The amino acid substitution Glu *β*39 → Arg, unique to crocodilian Hbs, appears to favour the R conformation over R2.

## 1 Introduction

Hemoglobins (Hbs) from all jawed vertebrates form similar hetero-tetramers, with two *α* and two *β* subunits ^1,2,3^. Hb is present at enormous concentration within the red blood cell, over 300 mg/mL in the case of humans for example, to facilitate oxygen transport around the body. Hb is one of the most studied proteins, and one of the first protein structures to be solved by X-ray crystallography. The Perutz group found that deoxy-Hb and oxy-Hb have different quaternary structures, which they associated with the T and R states of the MWC allosteric model ^4,5^. The change between these two quaternary structures of Hb can be described approximately as a rigid rotation of one *αβ* dimer relative to the other by about 15°, giving a different pattern of hydrogen bond interactions between the subunits. The MWC model explains the cooperative binding of oxygen by low affinity of the T state, which is stabilized by intersubunit bonds that oppose oxygen binding, so that the less contrained R state shows higher ligand affinity and is preferred at higher oxygen saturation. This model places all control of ligand affinity on the allosteric balance between T and R forms of different but invariant affinity. A novel quaternary structure of liganded HbA was identified in 1992 by the group of Arnone ^6^ and named R2. This model shows a greater dimer rotation from the T-state than the earlier crystallographic models, triggering extensive discussion of the actual conformation of the liganded molecule *in vivo*.

Many animal Hbs respond to small molecules within the red cell such as diphosphoglycerate (DPG), which selectively bind to T-state Hb and stabilize this form, reducing oxygen affinity ^7,8^. Adult human Hb (HbA) also binds carbon dioxide (largely present in the blood as bicarbonate ions) more strongly in the deoxy form, but this so-called Haldane effect is much stronger in crocodilian Hb, and independent of pH ^9,10,11,12^. In the deoxy state, crocodilian Hbs bind two bicarbonate ions per tetramer, with a dissociation constant close to 2 mM, but the oxygenated protein shows no appreciable bicarbonate binding. Bicarbonate ions therefore reduce oxygen affinity strongly, increasing the oxygen pressure at half-saturation (p50) of crocodilian blood by 40 mmHg and reducing the Hill cooperativity coefficient. Many vertebrate Hbs, including human and crocodile Hbs, have 141 residues in the *α* subunits and 146 residues in the *β* subunits. Helical regions are labelled from A to H ^13,14^. Hb from the American alligator *Alligator mississipiensis* (HbAM) shows 68% sequence identity to HbA in the *α* subunits, and 58% in the *β* subunits. The *α*_1_*β*_1_ and *α*_2_*β*_2_ interfaces are relatively fixed on allosteric transition of the protein, but the interfaces between these dimers show significant differences, especially at the so-called “switch” interface between the *α* subunit C helix and the *β* subunit FG corner ^15,16^.

With a molecular weight of around 64 kDa, Hb remains a small protein for cryo-EM analysis, but recent improvements in hardware and software have made it possible. Models of oxidized (met) Hb have been determined by cryo-EM using a Volta phase plate ^17^ or ultraflat graphene-supported grids ^18^. Higher resolution (2.8 Å) was achieved with more conventional methods by Herzik and colleagues ^19^. All of these studies used commercially available lyophilized human metHb, which may have been a factor in limiting resolution. The Herzik group found that *αβ* dimers accounted for about 20% of their data, which gives some indication of the sample quality ^19^. Instead, we have recently used conventional cryo-EM single particle analysis of fresh samples of human and crocodilian Hb, stored under carbon monoxide to prevent oxidation ^20^. That study focused on the allosteric mechanism of crocodilian Hbs, and the control of the allosteric equilibrium by bicarbonate ions. In this paper it has been attempted to provide a more thorough investigation of conformational space of liganded Hb by re-analyzing our previously published data-sets for the carbonmonoxy Hb samples. Models of human HbA in both R and R2 conformations are described from the same data-set. Moreover, the analysis of HbAM uncovered a previously uncharacterized conformation, which we term “T-like”, but not the R2 form, indicating a very different conformational equilibrium in the two proteins.

## 2 Results

Datasets for human Hb (HbA) in carbonmonoxy form (EMPIAR-11991) and HbAM in carbonmonoxy form (EMPIAR-11988) were described earlier ^20^. The data processing flow for re-analysis of each data-set is shown in Figs. 1 and 2; Fourier Shell Correlation curves for each map are shown in Fig. S1. Roughly 1.2 million particles of CO-bound human HbA and 900,000 particles of CO-bound HbAM were subjected to non-uniform consensus refinement with C2 symmetry for particle alignment. For HbA, the particles were then classified into eight classes using a mask covering the entire tetrameric map by three-dimensional (3D) classification with cryoSPARC ^21^, with classes labeled 1-8 based on particle count. The particle count and ratio are shown in Fig. 1. Non-uniform refinement was performed for all eight classes. Our previously described CO-bound human HbA model ^20^ (PDB:8WJ0) was used as a starting model, and all subunits were individually fitted into each reconstructed map using ChimeraX ^22^. Classes 1, 3, and 5 showed an *αβ* dimer rotation angle (compared to the deoxy model PDB:2DN2) of 20-22°, close to that of the R2-state conformation. The map of class 1 has the highest resolution, and its model has a 22° rotation. Class 2 showed a rotation angle of 16.3°, consistent with the R conformation. Classes 3 and 5 showed a rotation angle of approximately 20°, and classes 6-8 showed a rotation angle of approximately 19°, potentially representing intermediate conformations or average maps generated from both R-like and R2-like structures. Class 4 gave a low-resolution map and was discarded; possibly this class became a general bin into which any otherwise unclassified particles were assigned. Additional map refinement was carried out separately for class 1 and class 2, and with two groups of less-populated classes, 3+5, and 6+7+8. For classes 1 and 2, one round of heterogeneous refinement was performed, followed by one round of non-uniform refinement. The maps for classes 1 and 2 were obtained from 148,562 and 123,614 particles respectively, giving resolution of 2.26 Å and 2.45 Å (Fig. 1 and S1a-d). For classes 3+5, a 2.86 Å map was obtained from 176,209 particles, and this intermediate form between R and R2 was named “R2_1_”. For classes 6+7+8, a 2.71 Å map of a slightly different intermediate (“R2_2_”) was obtained from 259,342 particles (Figs. 1 and S1e-h). Model building was carried out with Coot ^23^, ISOLDE ^24^ and Servalcat ^25^. The models derived from the R2_1_ and R2_2_ maps are less reliable than the other models, and are not discussed in detail. Neither of them has been deposited in PDB. Data collection and refinement statistics for all the models, and database indentifier codes for the deposited models, are given in Table 1.

**Figure 1:**
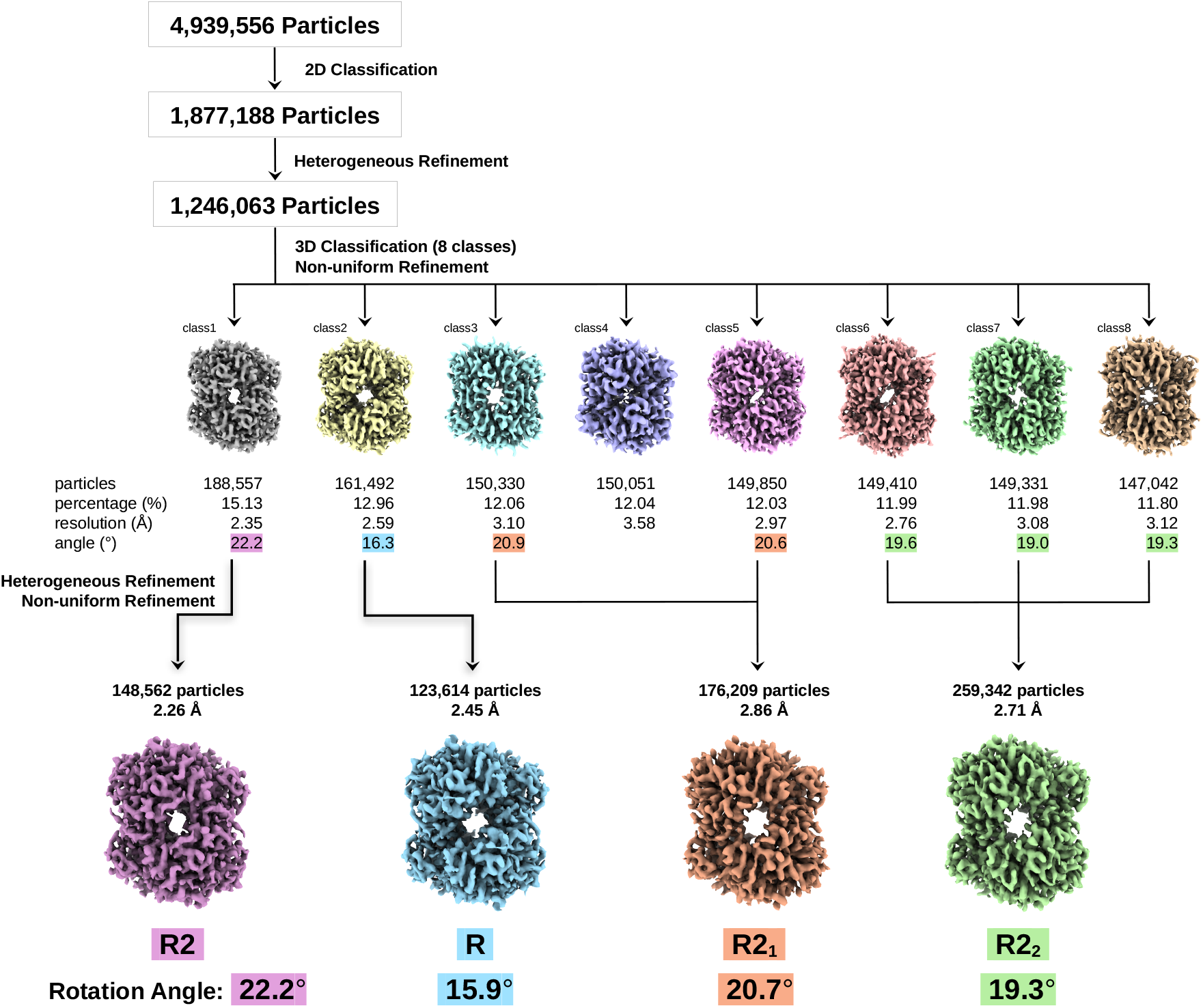
The workflow of the data processing and 3D classification analysis for CO-bound human HbA. Eight classes of particle were identified, and the corresponding maps are each shown with a unique colour. The population of each class relative to the identified particles is given as a percentage, together with the apparent angle of the *αβ* dimer rotation relative to deoxy-HbA (PDB:2DN2). The four final refined maps are shown, with arrows indicating the classes that contributed to each. The particle count is shown for each map, together with its resolution.

**Figure 2:**
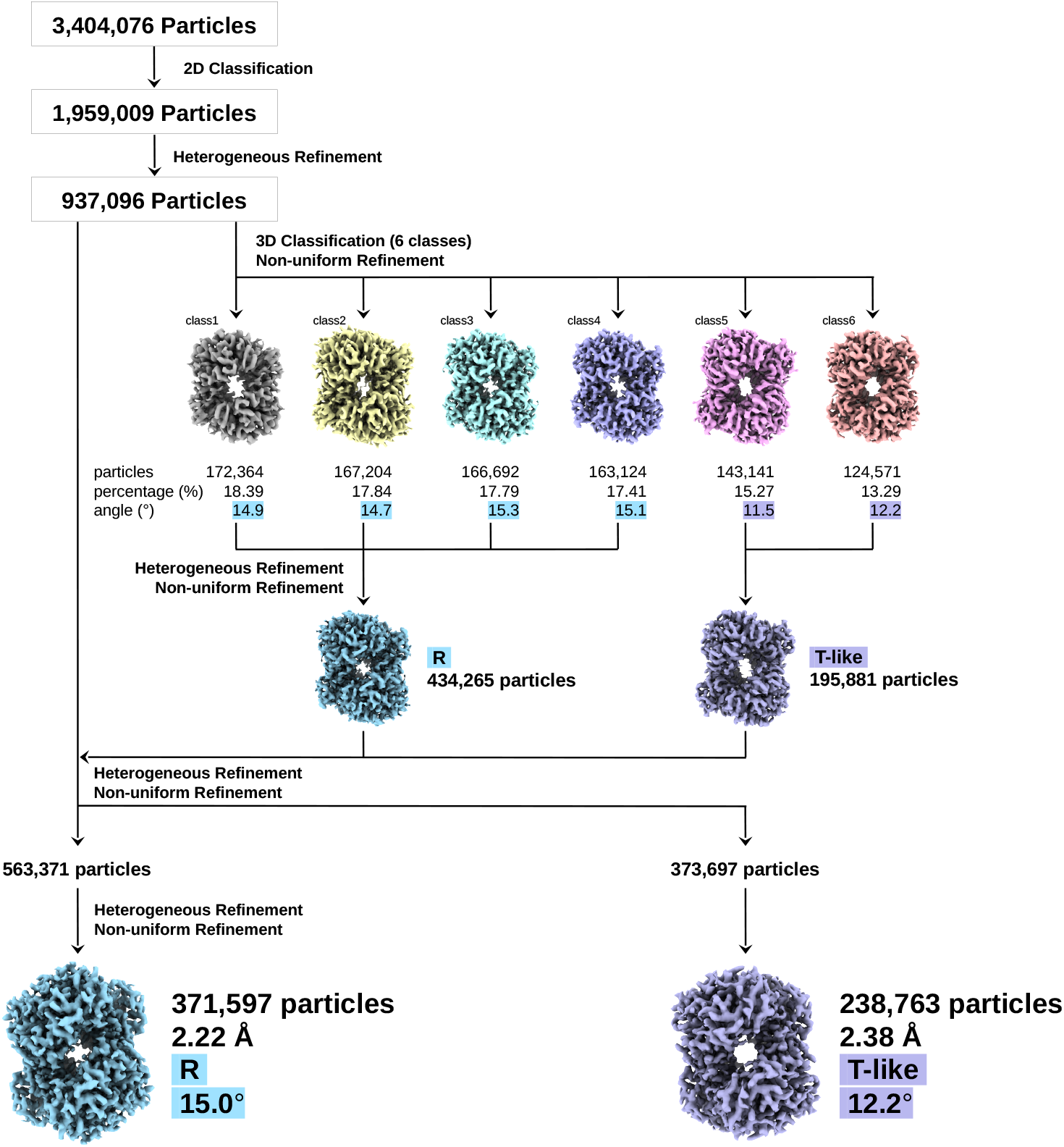
The workflow of the data processing and 3D classification analysis for CO-bound Hb from *Alligator missis-sipiensis*. Six classes of particle were identified, and the maps are each shown with a unique color. Similar data are presented as for the HbA models shown in Figure 1. The final maps of the R and crocodile-specific T-like structure are shown, coloured blue and purple respectively.

**Table 1.**
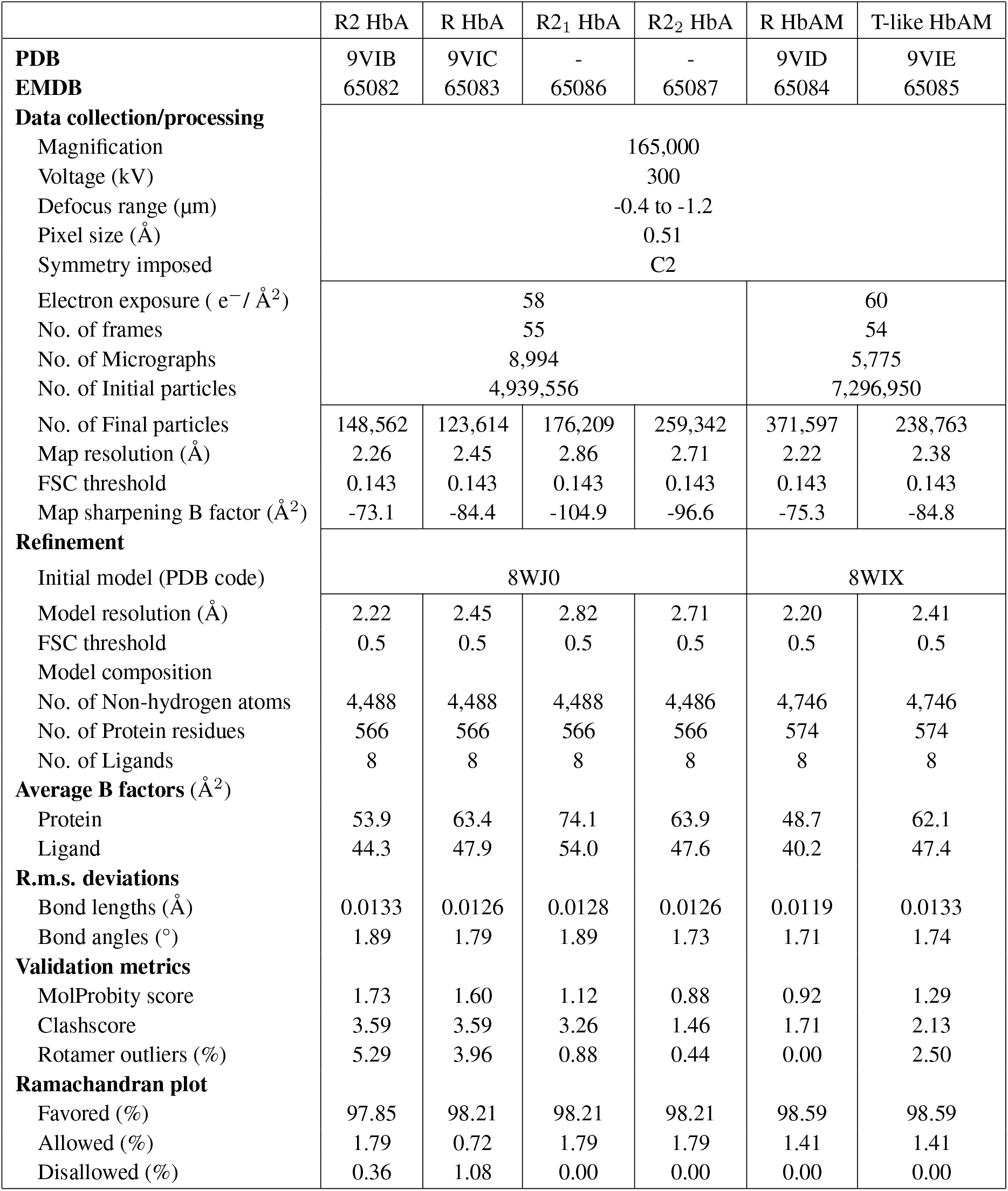
Data collection and refinement statistics.

**Table 2.** Atomic distances in various HbA and HbAM models. The columns R and HbAM(T-like) refer to the HbA and HbAM models described in this paper. 8WJ0 and 8WIX are HbA (R2) and HbAM (R) models in the PDB, described previously. Crosses indicate an atomic separation greater than 3.5 Å. 1BBB and 2DN3 are crystallographic models of CO-HbA in the R2 conformation and oxy-HbA in the R conformation respectively. The first three listed interactions indicate the T conformation, while the Asp *α*94-Asn *β*102 hydrogen bond is a key indicator of the R-state. Arg *β*39 and Tyr *β*41 are residues unique to crocodilian Hbs. Additional comparison between R and T conformation models is shown in Tables S5 and S6.

|  |  | R | 8WJ0 | 1BBB | 2DN3 | 8WIX | HbAM(T-like) | 8WIZ |
| --- | --- | --- | --- | --- | --- | --- | --- | --- |
| Lys $\alpha_1$127 Nζ | Arg $\alpha_2$141 O | × | × | × | × | × | × | 2.62 |
| Lys $\alpha_1$40 Nζ | His$\beta_2$146 O | × | × | × | × | × | 3.20 | 2.72 |
| Tyr $\alpha_1$42 Oη | Asp $\beta_2$99 Oδ | × | × | × | × | × | 2.25 | 2.48 |
| Asp $\alpha_1$94 Oδ | Asn $\beta_2$102 Nδ | 3.16 | 2.96 | 2.75/2.77 | 2.74 | 2.93 | × | × |
| Asn $\alpha_1$97 Nδ | Asn $\beta_2$99 Oδ | × | × | × | × | × | 3.05 | 2.94 |
| Arg $\alpha_1$141 O | Arg$\beta_2$39 Nη | - | - | - | - | 3.43/3.47 | × | × |
| Arg $\alpha_1$94 Oδ | Tyr$\beta_2$41 Oη | - | - | - | - | 2.43 | × | × |

The final class 1 model showed a dimer rotation angle of 22.2°, and represents an R2 conformation essentially the same as the previously published model (PDB:8WJ0). The class 2 model is more similar to the R conformation, with a dimer rotation angle of 15.8°, and is referred to in this paper as “R”. Some N-and C-terminal residues of liganded Hb models are not reliably positioned due to flexibility, and are best omitted from superposition calculations. Comparing the C*α* atoms of 564 residues (*α* subunit residues 2-141, *β* subunit residues 2-143) of the cryoEM R model with oxy-HbA (PDB:2DN3) or R2 HbA (PDB:1BBB) gives RMSD values of 0.98 Å and 1.28 Å respectively. Comparing the same atom set in PDB:8WJ0 with PDB:2DN3 and PDB:1BBB gives RMSD values of 1.45 Å and 0.58 Å respectively. Atomic distances that indicate the quaternary structure are listed in Table 2. Complete lists of close atom pairs (oxygen or nitrogen) in the two *αβ* pairs of the cryo-EM R and R2 models of human CO-Hb are given in Tables S1 and S2.

The interactions found at the *α*_1_*β*_2_ subunit interface of the new R conformation model of human CO-HbA refined in this work (class 2) are essentially identical to those of the X-ray crystallographic R structure of the same protein ^26^. In both the R and R2 conformations, the C helix of the *α*_1_ subunit approaches the C helix of the *β*_2_ unit (Fig. 3). Arg *β*40 can form a hydrogen bond with the Thr *α*41 side-chain of R-form HbA (which is replaced with isoleucine in HbAM). The interactions between the *α*_1_ subunit (Thr *α*41 and Tyr *α*42) and the Arg *β*_2_40 side-chain, and between side-chains of Asp *α*_1_94 and Asn *β*_2_102, are also found in the new HbA R model described here (Fig. 3c,f). The R and R2 conformations of Hb are characterized by different positions of the penultimate *α*_1_ subunit residue with respect to the *β*_2_ subunit (Fig. 4b-d and Table S3). In the R structure (but not R2) Pro *α*_1_95 and Tyr *α*_1_140 lie close to the highly conserved Trp *β*_2_37. Tyr *α*140 is clearly visible in the class 2 map in the R position, forming a hydrogen bond with the carbonyl oxygen of Val *α*93 (Fig. 4c). In the cryoEM R2 model and PDB:1BBB, Tyr *α*_1_140 forms a hydrophobic interaction with Val *α*_2_1 instead of Trp *β*_2_37 (Fig. 4b). Our previously described human HbA cryoEM model (PDB:WJ0) with the R2 conformation shows very similar interactions between the *α*_1_ and *β*_2_ subunits as the crystal structure PDB:1BBB (Fig. 3b,e). The interactions formed by Arg *α*141 in the R2 and R2-like structures are shown in Fig. S2.

**Figure 3:**
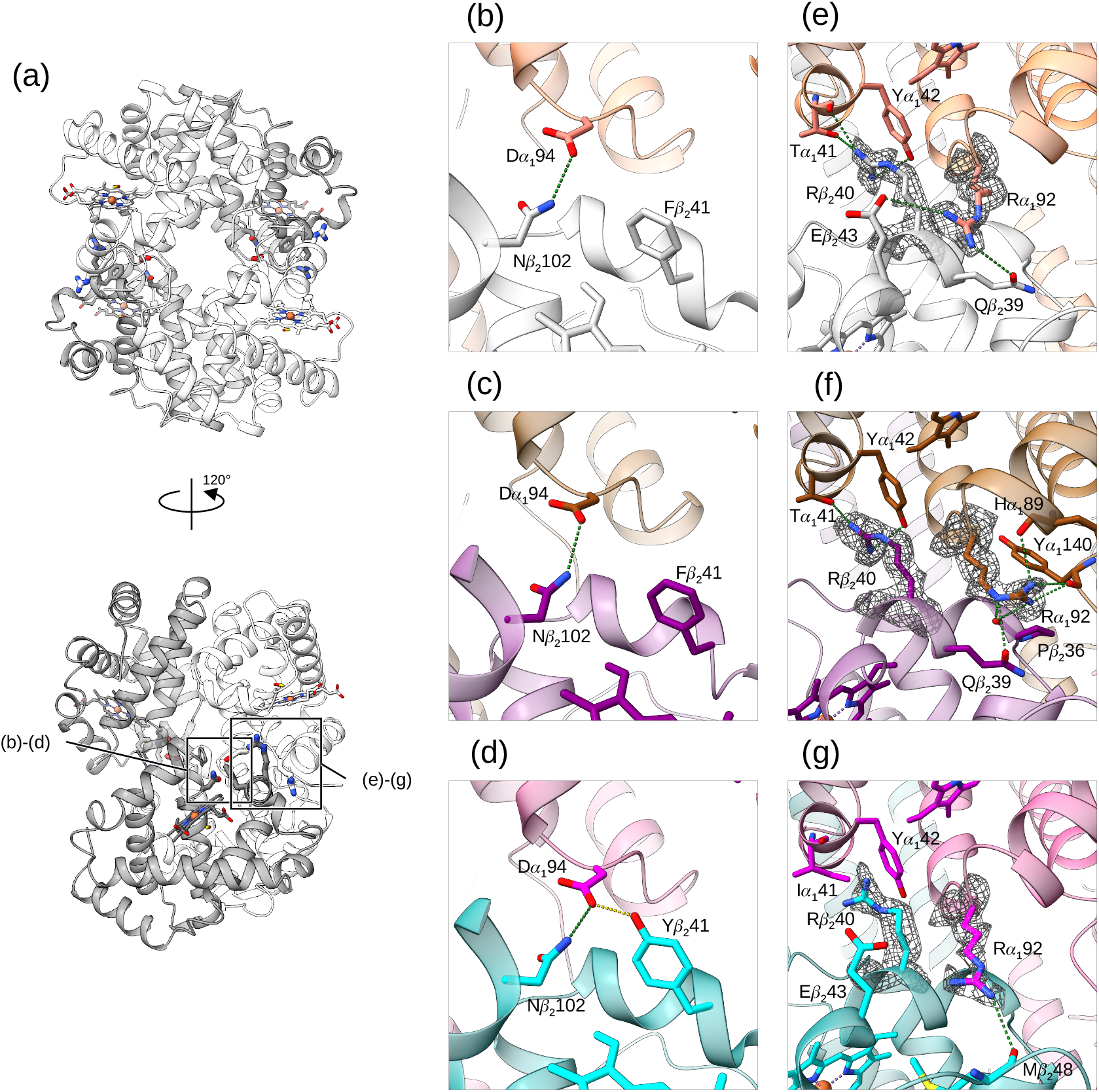
Interactions around the *α*_1_*β*_2_ interface. (a) Two different views of the cryo-EM R2 human HbA structure, shown as ribbon models. Boxes indicate the regions highlighted in panels (b) to (g). The carbon atoms of the *α* and *β* subunits are shown in white and grey respectively. The molecules of carbon monoxide are shown as sticks with carbon atoms coloured yellow. (b-d) The interactions formed by Asp *α*_1_94 in the R2 and R conformations, and the R-structure of HbAM, respectively. (e-g) Models and map density covering Arg *α*_1_92 and Arg *β*_2_40 at the *αβ* interface of R2 HbA, R HbA, and R HbAM, respectively. The carbon atoms of the *α* subunits of the R2-structure human HbA are shown in salmon, and the *β* subunits in white (b, e). The carbon atoms of the *α* subunits of the R-structure human HbA are shown in brown, and the *β* subunits in purple (c, f). The carbon atoms of the *α* subunits of the R2-structure HbAM are shown in magenta, and the *β* subunits in cyan (d, g). Atomic distances under 3.5 Å are shown as green dots, and the crocodile specific interaction (between Asp *α*_1_94 and Tyr *β*_2_41) is shown in gold (d).

**Figure 4:**
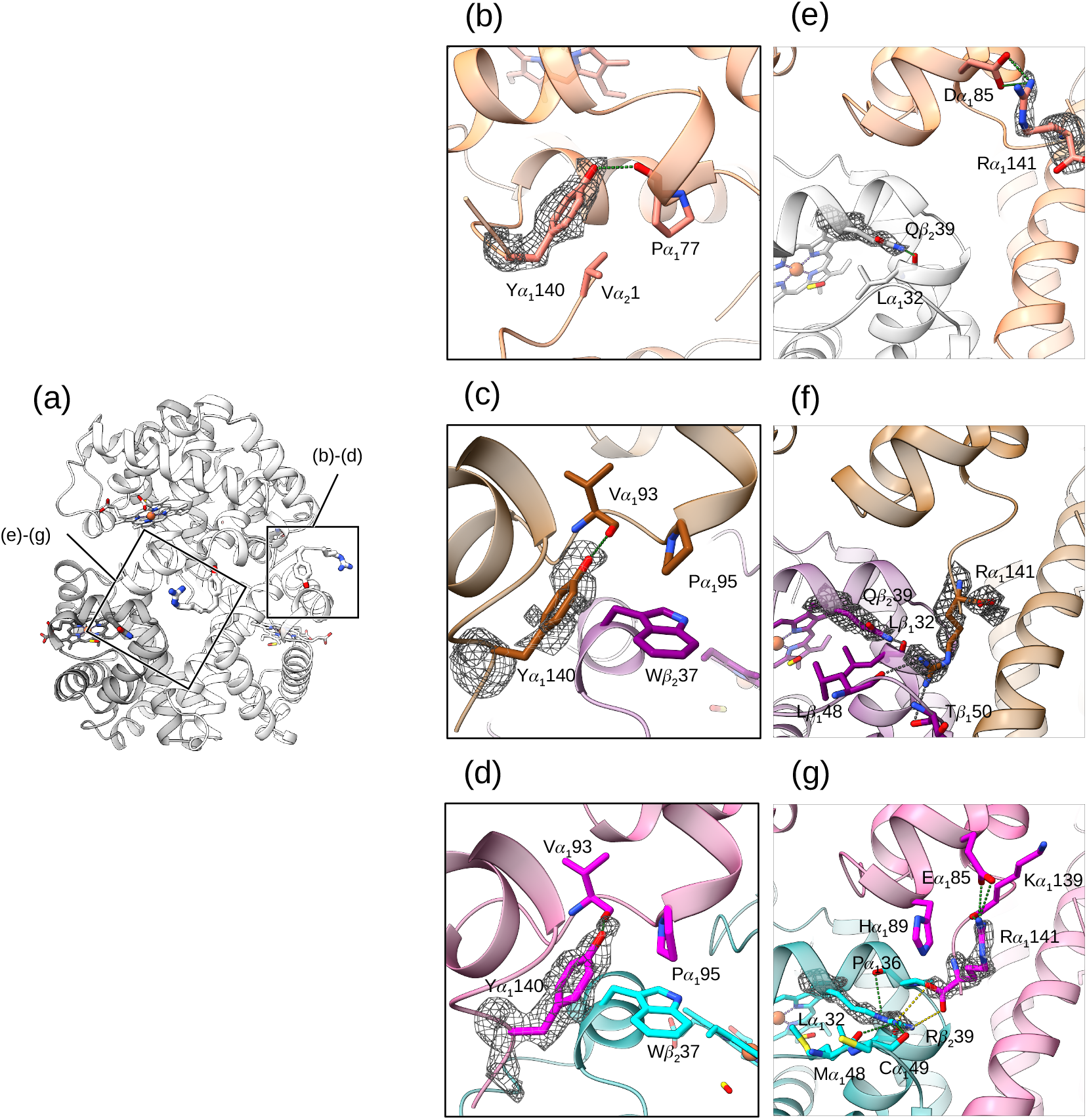
Interactions around the *α* subunit C-terminus. Intersubunit interactions around the C-terminus of the *α*_1_ subunit. Models are coloured as described in Figure 2. (a) The side view of the cryo-EM map of the R2-structure human HbA, with boxes showing the location of the highlighted features in panels b-g. Carbon monoxide ligands are shown as stick models (carbon atoms in yellow). (b-d) The environment of the Tyr *α*_1_141 subunit residue of (b) the R2 and (c) R structures of human HbA, and (d) the same region of R conformation HbAM. The map covering the tyrosine side-chain is shown. (e,f,g) The interactions between the C-terminus of the *α*_1_ subunit and residue *β*_2_39 of the R2 and R models of HbA, and the R-conformation model of HbAM, respectively. The map is shown over Arg *α*_1_141 and Gln/Arg *β*_2_39. Close atomic distances are shown as green dots, and the crocodile specific interaction formed by Arg *β*_2_39 and the *α* subunit C-terminus are shown in gold (g).

For HbAM, the workflow and data processing statistics are shown in Fig. 2. The particles were classified into six classes using a mask covering the entire tetrameric map, with classes labelled 1 to 6 based on particle count. Non-uniform refinement was performed for all six classes. Our previous CO-bound HbAM model ^20^ determined using cryo-EM (PDB:8WIX) was used as a reference model and fitted into each reconstructed map. The particles from classes 1 to 4 were combined for one round of heterogeneous refinement, followed by non-uniform refinement of the resulting major class. A map with resolution 2.33 Å was obtained from 434,265 particles, and the refined model shows a dimer rotation angle of 15.1°. This model is essentially identical to the one reported earlier (PDB:8WIX), and was not further analyzed. Classes 5 and 6 led to a 2.53 Å resolution model with a dimer rotation angle of about 12°. Since these two refined maps are clearly different from each other, one round of heterogeneous refinement was performed to classify the initial 937,096 particles using the two maps for reference. After one round of non-uniform refinement for each class, an R conformation map was produced from 563,371 particles, and a T-like map from 373,697 particles. One more round of heterogeneous refinement and non-uniform refinement for each class produced the final maps for R (371,597 particles and 2.22 Å resolution) and T-like forms (238,763 particles and 2.38 Å resolution). The rotation is overall in a direction opposite to the expected T-to-R transition, resulting in T-like features, such as hydrogen bonds between Tyr *α*42 and Asp *β*_2_99, and Lys *α*40 and His *β*_2_146 (Table 2 and Fig. S3). These interactions are also found in deoxy (T) HbAM (PDB:8WIZ). The Arg *β*40 side-chain and Arg *α*92 carbonyl oxygen form part of the bicarbonate ion binding site in PDB:8WIZ, but in the absence of this heterotopic effector these residues may adopt different conformations. Substantial disruption of the bicarbonate ion binding site is seen in the T-like structure, which prevents the ligand from binding. Instead of interacting with Glu *β*_2_43, the Arg *α*_1_92 side-chain lies against Trp *β*_2_37, with the indole ring adopting a flipped rotamer. The *αβ* dimer rotation also moves Tyr *α*_1_42 away from Tyr *β*_2_41 by almost 1 Å, preventing these two side-chains from both hydrogen-bonding effectively to a bicarbonate ion.

Hydrogen bonds between oxygen or nitrogen atoms in the refined CO-liganded HbAM models are shown in Tables S4 and S5. Bonds characteristic of the R conformation (such as between Asp *α*_1_94 and Asn *β*_2_102) are found in the new HbAM R model, but not in the T-like conformation. Unlike human HbA, therefore, liganded HbAM does not adopt an R2-like conformation, but does show a proportion in a novel conformation, with RMSD values of 1.98 Å, 3.10 Å and 4.37 Å when compared with PDB:2DN2 (deoxy HbA), PDB:2DN3 (oxy HbA) and PDB:1BBB (carbonmonoxy R2 HbA) respectively. Against the same three PDB models of HbA, PDB:8WIX shows RMSD values of 2.69 Å, 1.08 Å and 1.65 Å respectively. The N-termini of the *β* subunits in the T-like conformation lie much closer together (about 18 Å) than other liganded Hb models as a result of the unusual dimer rotation, which moves the centre-of-mass of the dimer in a very different direction from the expected T-to-R motion (Fig. S4), and leaves several T-marker hydrogen bonds intact (Table S6).

The interaction between Asp *α*94 and Asn *β*102 side-chains is preserved in both human HbA and HbAM, in the R and R2 conformations (Fig. 3e-g). In the HbAM R model, Asp *α*94 also interacts with Tyr *β*41, a residue unique to crocodilian Hbs (Fig. 3g). This interaction is not found in human HbA, where residue *β*41 is replaced by the generally conserved phenylalanine. Interactions around the *α* subunit C-terminus are shown in Fig. 4. Arg *α*141 of HbA lies away from Gln *β*_2_39 in the R2 conformation of HbA (Fig. 4b), but much closer in the R conformation (Fig. 4c). Gln *β*39 can form a hydrogen bond with the carbonyl oxygen of Leu *β*32. In liganded HbAM (PDB:8WIX), the replacement of Gln *β*39 with arginine allows this reside to form a salt bridge with the C-terminal carboxyl group of Arg *α*_2_141. The Arg *α*141 side-chain forms a cation-*π* interaction with His *α*89 and a hydrogen bond with the carbonyl oxygen of Lys *α*139 (Fig. 4d). In the R and R2 conformations of HbA, Gln *β*39 can form a hydrogen bond with the carbonyl oxygen of Leu *β*32. In the R2 form, the *α*_1_ C-terminal residue Arg *α*141 and Gln *β*_2_39 of HbA move apart so that Arg *α*141 does not form any interactions with the *β* subunit, and the side-chain forms a salt bridge with Asp *α*85 (Fig. 4b). A model of the R2-state of HbAM was built by fitting each HbAM subunit to the EM-map of the R2-state human HbA.The subunit interactions in the R2 human HbA model PDB:1BBB are largely conserved, but the Arg *β*39-Arg *α*141 salt-bridge would be broken in the R2 conformation.

## 3 Discussion

The earliest crystallographic studies of Hb by Perutz, using high concentrations of salts, produced models of both liganded and unliganded Hb, but offered little insight into the conformational variation within each of these different states, although it was appreciated that salt bridges in the T-state could be broken without a complete switch to the R-state. A great deal of debate was triggered in the 1990s by the structure of a functionally impaired mutant, Hb Ypsilanti (Asp *β*99→Tyr), which has a destabilized T-state ^27^. It was suggested that this mutant reflects a third quaternary state of normal HbA. Much stronger evidence for a biologically important novel liganded conformation came from the group of Arnone, which grew crystals of carbonmonoxy human HbA (CO-HbA) using polyethylene glycol (PEG) as precipitant, and relatively low ionic strength ^6^. The principal difference between the earlier “high-salt” oxyHb crystal structure (PDB:1HHO) ^28^ and the Arnone structure (PDB:1BBB) is a largely a simple rotation of one rigid *αβ* pair relative to the other, much in the way that the Perutzian T and R structures differ. The new quaternary structure was named R2. The rotation from T to the R2 form is greater (and with a slightly different axis) than the rotation to the R state. It was therefore suggested that the R2 state is a “hyper-R state”, that lies beyond the T-to-R transition, and furthermore that the Arnone model was more likely to represent the structure of the protein in solution; it was proposed that the “high-salt” crystallization conditions used by Perutz impeded the full dimer rotation, creating an artefact ^29^. No evidence emerged to support the proposal that high ionic strength could have this effect, and protein crystal structures solved using salt precipitants appear generally to reflect well structural models of the same protein obtained under different conditions, or by other methods. Very high resolution crystal structures of HbA in different ligation states later showed that the oxyHb model PDB:1HHO has a peptide flip error which previous analyses had taken to be diagnostic of the R conformation ^26^. The crystals used to determine the R2 model were grown at relatively low pH (5.8) ^6^, and there was therefore great interest in a subsequent report of crystals of cyanomet HbA, grown in PEG at pH 7.4^30^, also suggested to show the R2 form, but the paper described only the result of a molecular replacement search. Almost 30 years later, no refinement has been reported, and no models or data have been deposited in a public database. Attempts to recover the data have proved fruitless, and there has been no report of any replication of these crystals. Nevertheless, at the time, this report provided a considerable boost to the then growing narrative that the R state models such as PDB:1HHO ^28^ are artefactual. To counter this argument, one of us (JRHT) wrote a short commentary in 1999, pointing out that by this time a number of other vertebrate Hbs had been solved by X-ray crystallography, using either high or low ionic strength crystallization conditions, all of them between pH 6.7 and 8.0^31^. Of 12 then available models (some of them cross-linked HbA), three liganded fish Hbs and goose Hb crystallized using PEG all showed conformations closer to R than R2. All six animal Hbs were more R-like, while only two were crystallized using high ionic strength. The purpose of the commentary was to provide an overview of all the evidence at the time, which did not strongly indicate the R conformation to be an artefact, and to point out that if the R and R2 structures have identical oxygen binding properties then distinguishing them becomes something of a semantic problem, as each structure may represent a point within the conformational space of the thermodynamic R state. The concluding paragraph included the sentence

> The structures of animal haemoglobins, however, seem to provide strong evidence that the R state is a better representation of the oxy-Hb molecule.

but suggested debate would continue and further studies were needed to resolve the R vs. R2 question. This drew a strongly critical response from Arnone and colleagues in an appendix to a paper describing three novel crystal structures of carbonmonoxy bovine Hb ^32^. These models, all crystallized under low ionic strength, showed various degrees of dimer rotation, prompting the authors to describe the inter-dimer contacts as a “sliding interface”. The paper revealed differences in views of what a protein structure represents, so that the Arnone group coined the term “R^*e*^” to indicate an ensemble of liganded structures at equilibrium in solution. It is notable however that all of the bovine structures reported show the same pattern of hydrogen bonds identified by Perutz decades earlier as characterizing the R-state ^5^. Identifying the quaternary structure of the protein through subunit interactions is certainly simpler (and possibly less arbitrary) than matching dimer rotation angles, as each hydrogen bond pattern maps onto substantial volumes of conformational space ^26^. Crystallographic studies suggest the ensemble formed by T-state Hb is more tightly distributed than that of R-state Hb, but nevertheless crystallization is a selective process so that each crystal structure may sample only a portion of the conformational space explored by the protein free in solution. The collapse of all conformational possibilities to a single-valued model for each *x, y* and *z* coordinate is not only due to this selection, but also the averaging of the crystallographic method, and the final model building. The quaternary flexibility of liganded Hb therefore presents an example where the ability to pick particles in cryo-EM images can provide deeper and more controlled analysis of conformational space than crystallography by a process of “on-grid purification” ^33^.

The discovery of different crystallization conditions using PEG has allowed studies of oxygenation intermediates, and even T-state HbA with four oxygen molecules bound with crystallographic methods ^34,35,36^. More recently, a crystal form has been described with three tetramers per asymmetric unit that can accommodate structures across the range from T to R2^37,38^. The T-and R-state conformations of this crystal form are consistent with a meta-analysis of many Hb structures in PDB ^39^. With the development of dipolar coupling methods, NMR became able to measure the average *αβ* dimer rotation of a liganded Hb sample in solution, showing that liganded HbA in solution has an average rotation angle roughly half-way between crystallographic R and R2 structures ^40,41^. Using cryo-EM it is possible to select individual particles, and average subsequently, which provides a much better means of exploring the conformational space of Hb. The analysis here shows that liganded human HbA in buffered saline solution exists in a variety of conformations, with R2 being the most prevalent. The R conformation is barely less populated, and two intermediate conformations are present in sufficient quantity for averaging and map refinement. In the case of crocodilian Hb, the R2 conformation is absent, and residue changes around the bicarbonate binding site strongly favour the R conformation over R2. This provides clear proof that the allosteric mechanism of Hb does not rely on the R2 conformation, merely breakage of intersubunit bonds specific to the T-state. Indeed, breakage of some bonds within the T-state can also increase oxygen affinity without a switch to the R conformation ^38^. The range of oxygen affinity expressed by different quaternary states of Hb under different solution conditions is considerable ^42,43^. Much confusion in this area has arisen from the loose approximation of a Hb conformation to a thermodynamic state, despite the clear fact that the average conformation may remain unchanged with a change in solution conditions (temperature, ionic strength and so on), while the thermodynamic state may change markedly in energy. Arg *β*39, associated with the bicarbonate binding effect of crocodilian Hbs, appears to be largely responsible for the preference of liganded HbAM to adopt the R conformation. The unexpected T-like structure has no obvious functional role, and appears unable to bind bicarbonate ions due to side-chain movements around the trio of positively-charged side-chains in the C helix of crocodilian *β* globins. Possibly this model reflects the disruption caused by the Lys *β*38 side-chain, which is crucial for bicarbonate binding but which displaces the highly conserved Trp *β*37^20^. More speculatively, the local disturbance and disorder arising at this site due to evolution of bicarbonate sensitivity may be responsible for the mysterious difficulty of crystallizing any crocodilian Hb in a form suitable for X-ray analysis. The Hb models presented here, as well as work from other groups cited above, suggest that human Hb may explore a larger conformational space in the liganded state than some other vertebrate Hbs. Whether this added entropic stabilization of the R-state is functionally important, and related to the high cooperativity (Hill coefficient) of HbA, remains to be tested. It is notable that wide-angle X-ray scattering (WAXS) studies of bovine carbonmonoxy-Hb suggest that this protein forms a structure between the T and R forms of HbA ^44^, which is consistent with the crystal structure (PDB:1FSX) refined by Safo and Abraham using 1.6 M phosphate as precipitant ^45^. Bovine Hb has an intrinsically low oxygen affinity, and in the presence of physiological chloride concentrations it does not respond to organic phosphates such as DPG, an effect that has been attributed to the replacement of the *β*2 side-chain with a large hydrophobic group ^8^. Human HbA has histidine at this position, and HbAM has serine. Bovine Hb has leucine instead, and cat Hb, with phenylalanine at the *β*2 position, also shows a very weak response to DPG, low intrinsic oxygen affinity, and low cooperativity ^46^. Crystal structures of oxidized (met) cat Hb such as PDB:3GQP show a mixture of T and R marker hydrogen bonds ^47^. The T-state marker hydrogen bonds of HbAM in the T-like form therefore suggest a reduced oxygen affinity compared to the R conformation, despite the high intrinsic oxygen affinity of HbAM in the absence of bicarbonate ions. WAXS studies of carbonmonoxy-HbA are inconsistent with any combination of R and R2 models ^44^, and the present cryo-EM models may go some way towards resolving the question of the structure of liganded human Hb in solution.

## 4 Materials and Methods

HbA was purified by Prof. Tsuneshige of Hosei University, Japan, using standard protocols and blood drawn from a human volunteer. The red cells were washed and bubbled with carbon monoxide before lysis, and the purified protein was shipped to Yokohama at 4^*°*^ under carbon monoxide in stoppered tubes. Carbonmonoxy-HbA was diluted to mg/mL with phosphate-buffered saline to prepare EM samples. 3 *µ*L drops of the protein were applied to grids (Quantifoil Cu/Rh R1.2/1.3 carbon holey 300 mesh), which were glow-discharged in advance (10 mA for 50 s), and blotted with a blot force of 10 at 6^*°*^ and 100% humidity with a Vitrobot (Mark IV, ThermoFisher), and then plunge frozen into liquid ethane.

Whole alligator blood was provided by Prof. Dane Crossley of The University of North Texas from nine adult alligators assigned to independent research projects. After euthanasia, blood was drawn using heparinized syringes according to UNT guidelines and stored at -80^*°*^. After thawing, red cells were collected by centrifugation at 4^*°*^, 4000 *× g*, and washed with ice-cold 0.9% sodium chloride containing 0.5 mM EDTA. The red cells were lysed by 1:5 dilution with ice-cold 10 mM HEPES pH 7.6, 0.5 mM EDTA, and incubated on ice for 45 min. The hemolysate was then centrifuged at 4^*°*^, 12000 × *g*, for 20 min. to remove cell debris. The hemolysate was passed through a PD-10 SephadexTM G-25 desalting column (GE Healthcare) previously equilibrated with 10 mM HEPES pH 7.6, 0.5 mM EDTA.

HbAM was purified by ion exchange chromatography using a HiTrap Q XL column (Cytiva, Uppsala) equilibrated with 20 mM HEPES pH 7.6, 0.5 mM EDTA, and then eluted with a linear 0-0.1 M NaCl gradient, at room temperature. The adult Hb fraction was concentrated by centrifugation, passed again through a PD-10 column, and dialysed against 10 mM HEPES pH 7.4, 0.5 mM EDTA to remove sodium chloride. CO-bound HbAM was diluted to a final concentration of 6.5 mg/ml with TN buffer (20 mM Tris and 100 mM NaCl pH 8.0) containing 10 mM dithiothreitol.

HbAM samples were applied to grids using the same protocol as HbA.

All cryo-EM data were collected at RIKEN Yokohama, using 300 kV acceleration voltage with a Titan Krios G4 (ThermoFisher) equipped with a K3 detector (Gatan) in counting mode with correlative double sampling (CDS), using automated EPU software. Images were acquired at a magnification of 160,000*×* and a pixel size of 0.51 Å. Movies were collected with 54-56 frames of -0.4 to -1.2 *µ*m defocus with two to three shots per hole. The total exposure time was 2.0 s, and the exposure dose was 7.6-7.8 e/pix/s; the total exposure was 58.4 - 60.0 e/Å^2^.

## Acknowledgement

We thank Profs. A. Tsuneshige and D. Crossley for their kind gifts of purified human HbA and alligator blood respectively. We thank Profs. Jay Storz and Angela Fago, and Dr Naim Bautista, for purifying the alligator Hb, and their continued support. KT acknowledges a JST university fellowship, grant number JPMJFS2140. We thank Profs. Beatrice Vallone and Naoya Shibayama for their comments on the manuscript.

## Data availability

The structural models have been deposited in the Protein Data Bank (PDB), and the cryo-EM density maps, half maps, and masks have been deposited in the Electron Microscopy Data Bank (EMDB) with following accession codes: human haemoglobin in the R2 conformation (9VIB, EMD-65082), human haemoglobin in the R conformation (9VIC, EMD-65083), human haemoglobin in intermediate conformation 1 (R2_1_) (EMD-65086), human haemoglobin in intermediate conformation 2 (R2_2_) (EMD-65087), alligator haemoglobin in the R conformation (9VID, EMD-65084), and alligator haemoglobin in the T-like conformation (9VIE, EMD-65085).

## Supplementary Information

**Figure S1:**
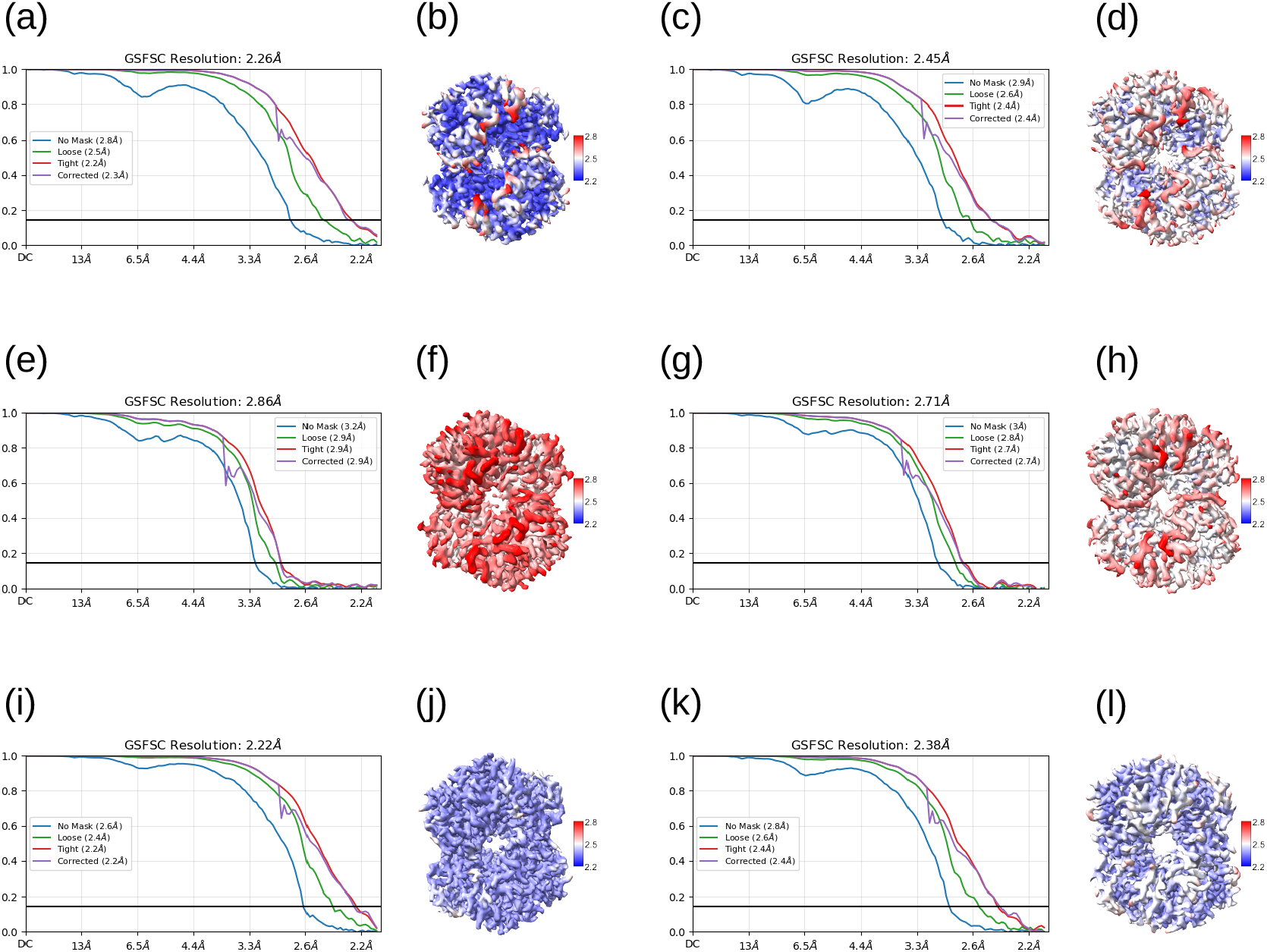
Interactions around the *α* subunit C-terminus. The Gold-Standard Fourier Shell Correlation (FSC) curves and the local resolution filtered EM maps coloured from high resolution in blue to low resolution in red. (a, b) R2 HbA. (c, d) R HbA. (e, f) R2_1_ HbA. (g, h) R2_2_ HbA. (i, j) R HbAM. (k, l) T-like HbAM.

**Figure S2:**
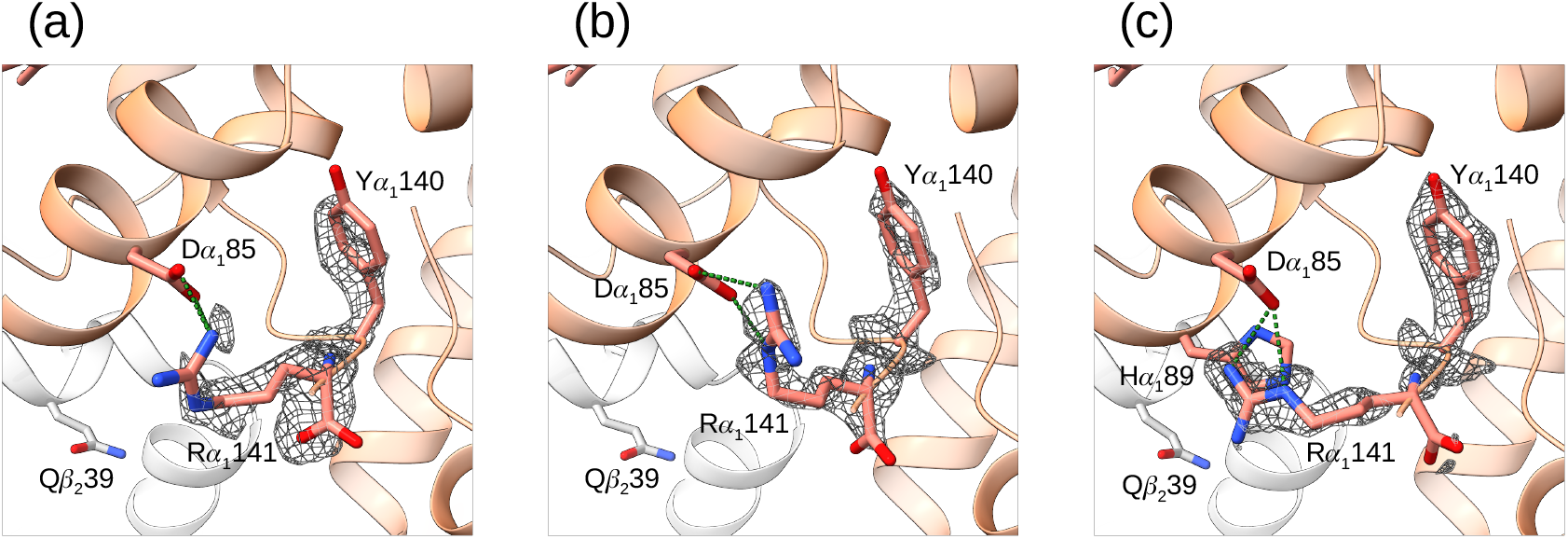
Interactions close to the *α* subunit C-terminus in the HbA R2 and R2-like intermediate structures. (a) R2 structure of human HbA. (b) R2_1_ structure. (c) R2_2_ structure. Colouring is the same as Figure 3. Close atomic distances are shown as green dots. The interaction between Arg *α*_1_141 and Gln *β*_2_39 is lost in all three of the models shown. Instead Arg *α*_1_141 approaches Asp *α*_1_85.

**Figure S3:**
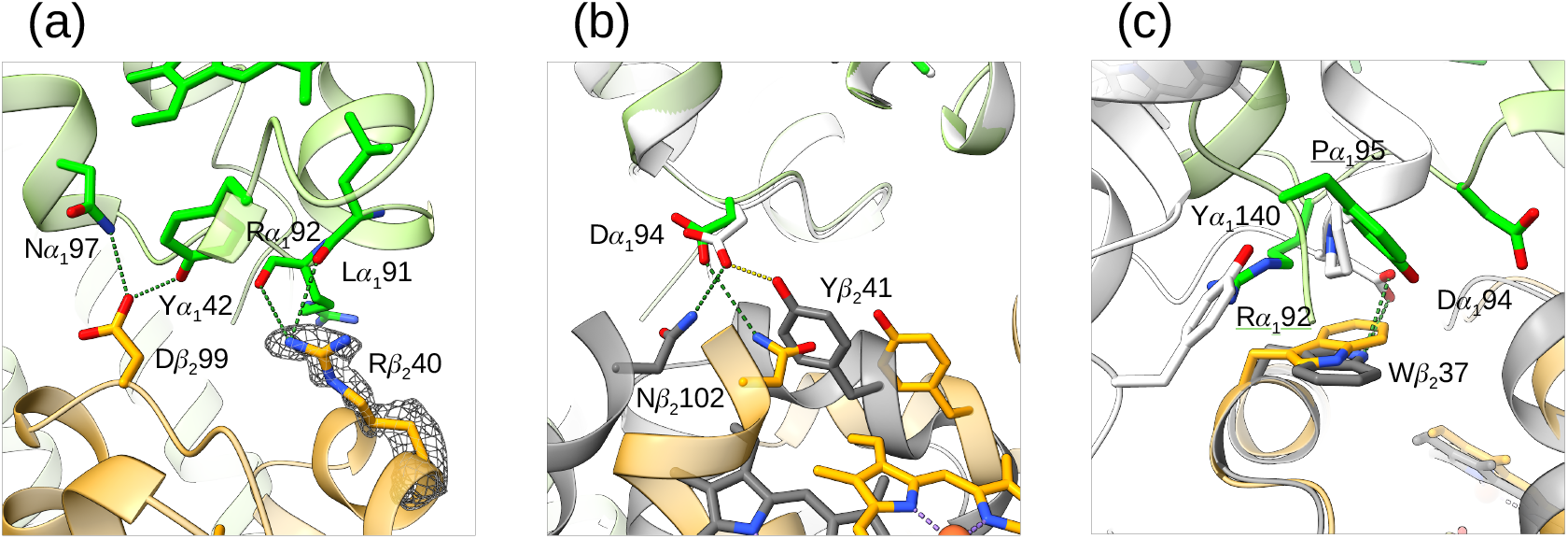
Subunit interactions at the *α*_1_*β*_2_ interface of T-like HbAM. (a) The carbon atoms of the *α* and *β* subunits of the T-like conformation HbAM model are shown in lime and orange respectively. Interactions formed by the *α*_1_subunit with Arg *β*_2_40 and Asp *β*_2_99. (b) The R model of HbAM is shown superposed on the T-like model by matching the *α*_1_ subunits. The subunits of the R-structure HbAM are shown in white (*α*) and grey (*β*). The two conformations show very different relative positions of Tyr *α*_1_94 and Tyr *β*_2_41. Close atomic distances are shown as yellow or green dots for the side-chains of Tyr *β*41 and Asn *β*102 respectively. (c) The same overlay as in (b), but showing the side-chain of Trp *β*37 has flipped rotamer in the T-like conformation.

**Figure S4:**
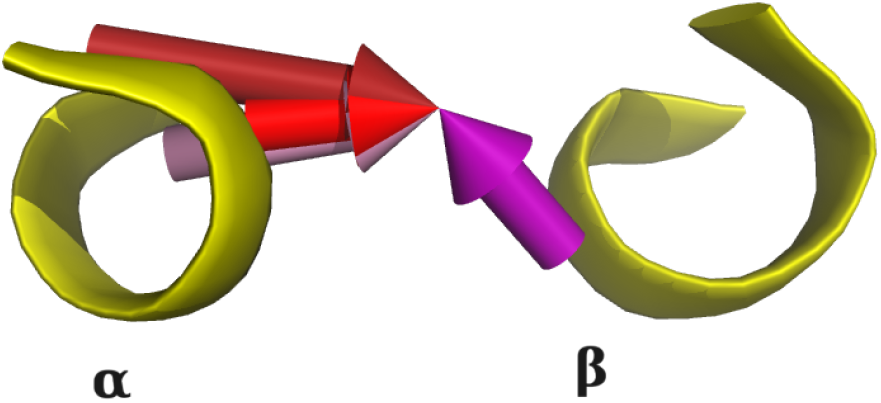
Movement of the centre-of-mass of the dimer rotation of different Hb models. Various liganded Hb models were overlaid onto deoxy HbA (PDB:2DN2) by fitting the *α*_1_*β*_1_ dimer, and then the *α*_2_*β*_2_ dimer was overlaid. Arrows representing the COM movement of this second fitting operation are shown, together with single turns from the G helices of the 2DN2 model *α*_2_ and *β*_2_ subunits. Arrows for oxyHbA (red), CO-HbA R2 (dark red), and HbAM R (pink) models all show the same general direction of movement, with R2 showing the largest change. T-like HbAM (purple) shows a very different motion, away from the classic T-to-R transition.

**Table S1:** Contacts in human carbonmonoxy HbA R conformation.

| Chain | Res | ResName | Atom | Chain | Res | ResName | Atom | Distance (Å) |
| --- | --- | --- | --- | --- | --- | --- | --- | --- |
| A | 41 | THR | OG1 | D | 40 | ARG | NH1 | 3.11 |
| A | 92 | ARG | NE | D | 36 | PRO | O | 3.29 |
| A | 92 | ARG | NE | D | 39 | GLN | OE1 | 3.35 |
| A | 92 | ARG | NH1 | D | 36 | PRO | O | 3.13 |
| A | 94 | ASP | OD2 | D | 102 | ASN | ND2 | 3.16 |
| A | 141 | ARG | NH1 | D | 50 | THR | O | 2.98 |
| A | 141 | ARG | NH1 | D | 51 | PRO | N | 2.98 |
| A | 141 | ARG | NH2 | D | 39 | GLN | NE2 | 3.27 |
| A | 141 | ARG | NH2 | D | 48 | LEU | O | 2.93 |
| B | 36 | PRO | O | C | 92 | ARG | NE | 3.29 |
| B | 36 | PRO | O | C | 92 | ARG | NH1 | 3.14 |
| B | 39 | GLN | OE1 | C | 92 | ARG | NE | 3.35 |
| B | 39 | GLN | NE2 | C | 141 | ARG | NH2 | 3.27 |
| B | 40 | ARG | NH1 | C | 41 | THR | OG1 | 3.11 |
| B | 48 | LEU | O | C | 141 | ARG | NH2 | 2.93 |
| B | 50 | THR | O | C | 141 | ARG | NH1 | 2.98 |
| B | 51 | PRO | N | C | 141 | ARG | NH1 | 2.98 |
| B | 102 | ASN | ND2 | C | 94 | ASP | OD2 | 3.16 |

Contacts between the $\alpha_1\beta_1$ and $\alpha_2\beta_2$ dimers, involving pairs of nitrogen or oxygen atoms, closer than 3.5 Å. All cryoEM models of Hb described in this paper have strict two-fold symmetry imposed in refinement.

**Table S2:** Contacts in human carbonmonoxy HbA R2 conformation.

| Chain | Res | ResName | Atom | Chain | Res | ResName | Atom | Distance (Å) |
| --- | --- | --- | --- | --- | --- | --- | --- | --- |
| A | 41 | THR | O | D | 40 | ARG | NH1 | 3.28 |
| A | 41 | THR | OG1 | D | 40 | ARG | NH1 | 3.42 |
| A | 94 | ASP | OD1 | D | 37 | TRP | NE1 | 3.47 |
| A | 94 | ASP | OD2 | D | 102 | ASN | ND2 | 3.29 |
| A | 127 | LYS | NZ | C | 139 | LYS | O | 3.12 |
| A | 139 | LYS | O | C | 127 | LYS | NZ | 3.12 |
| B | 37 | TRP | NE1 | C | 94 | ASP | OD1 | 3.47 |
| B | 40 | ARG | NH1 | C | 41 | THR | O | 3.28 |
| B | 40 | ARG | NH1 | C | 41 | THR | OG1 | 3.42 |
| B | 102 | ASN | ND2 | C | 94 | ASP | OD2 | 3.29 |

Contacts between the $\alpha_1\beta_1$ and $\alpha_2\beta_2$ dimers, involving pairs of nitrogen or oxygen atoms, closer than 3.5 Å.

**Table S3:** Distances between C*α* atoms in various liganded Hb models.

|  |  | R | R2 | 1BBB | 2DN3 | HbAM | HbAM(T-like) | 1FSX |
| --- | --- | --- | --- | --- | --- | --- | --- | --- |
| Tyr $\alpha_1$140 | Tyr $\beta_2$37 | 6.9 | 14.3 | 14.1/14.3 | 7.3 | 6.7 | 9.3 | 8.0/8.3 |
| Arg $\alpha_1$141 | Tyr $\beta_2$37 | 8.6 | 15.8 | 15.9/16.3 | 8.9 | 8.1 | 14.1 | 10.2/10.6 |
| $\beta_1$2 | $\beta_2$2 | 28.3 | 30.9 | 33.1 | 26.8 | 28.0 | 17.8 | 32.9 |

R refers to the CO-HbA model described in this paper, and R2 refers to PDB:1WJ0. PDB:1BBB and PDB:1FSX are models of CO-HbA in the R2 conformation, and bovine CO-Hb respectively.

**Table S4:** Contacts in alligator carbonmonoxy Hb (CO-HbAM) R conformation.

| Chain | Res | ResName | Atom | Chain | Res | ResName | Atom | Distance (Å) |
| --- | --- | --- | --- | --- | --- | --- | --- | --- |
| A | 92 | ARG | NH1 | D | 48 | MET | O | 3.43 |
| A | 94 | ASP | OD1 | D | 37 | TRP | NE1 | 3.47 |
| A | 94 | ASP | OD1 | D | 99 | ASP | OD2 | 3.47 |
| A | 94 | ASP | OD2 | D | 41 | TYR | OH | 2.39 |
| A | 94 | ASP | OD2 | D | 102 | ASN | ND2 | 2.85 |
| A | 141 | ARG | O | D | 39 | ARG | NH1 | 3.21 |
| B | 37 | TRP | NE1 | C | 94 | ASP | OD1 | 3.47 |
| B | 39 | ARG | NH1 | C | 141 | ARG | O | 3.21 |
| B | 41 | TYR | OH | C | 94 | ASP | OD2 | 2.39 |
| B | 48 | MET | O | C | 92 | ARG | NH1 | 3.43 |
| B | 99 | ASP | OD2 | C | 94 | ASP | OD1 | 3.47 |
| B | 102 | ASN | ND2 | C | 94 | ASP | OD2 | 2.85 |

Contacts between the $\alpha_1\beta_1$ and $\alpha_2\beta_2$ dimers, involving pairs of nitrogen or oxygen atoms, closer than 3.5 Å.

**Table S5:** Contacts in alligator carbonmonoxy Hb (CO-HbAM) T-like conformation.

| Chain | Res | ResName | Atom | Chain | Res | ResName | Atom | Distance (Å) |
| --- | --- | --- | --- | --- | --- | --- | --- | --- |
| A | 40 | LYS | NZ | D | 146 | HIS | OXT | 3.20 |
| A | 42 | TYR | OH | D | 99 | ASP | OD2 | 2.25 |
| A | 97 | ASN | ND2 | D | 99 | ASP | OD2 | 3.05 |
| B | 99 | ASP | OD2 | C | 42 | TYR | OH | 2.25 |
| B | 99 | ASP | OD2 | C | 97 | ASN | ND2 | 3.05 |
| B | 146 | HIS | OXT | C | 40 | LYS | NZ | 3.20 |

Contacts between the $\alpha_1\beta_1$ and $\alpha_2\beta_2$ dimers, involving pairs of nitrogen or oxygen atoms, closer than 3.5 Å

**Table S6:**
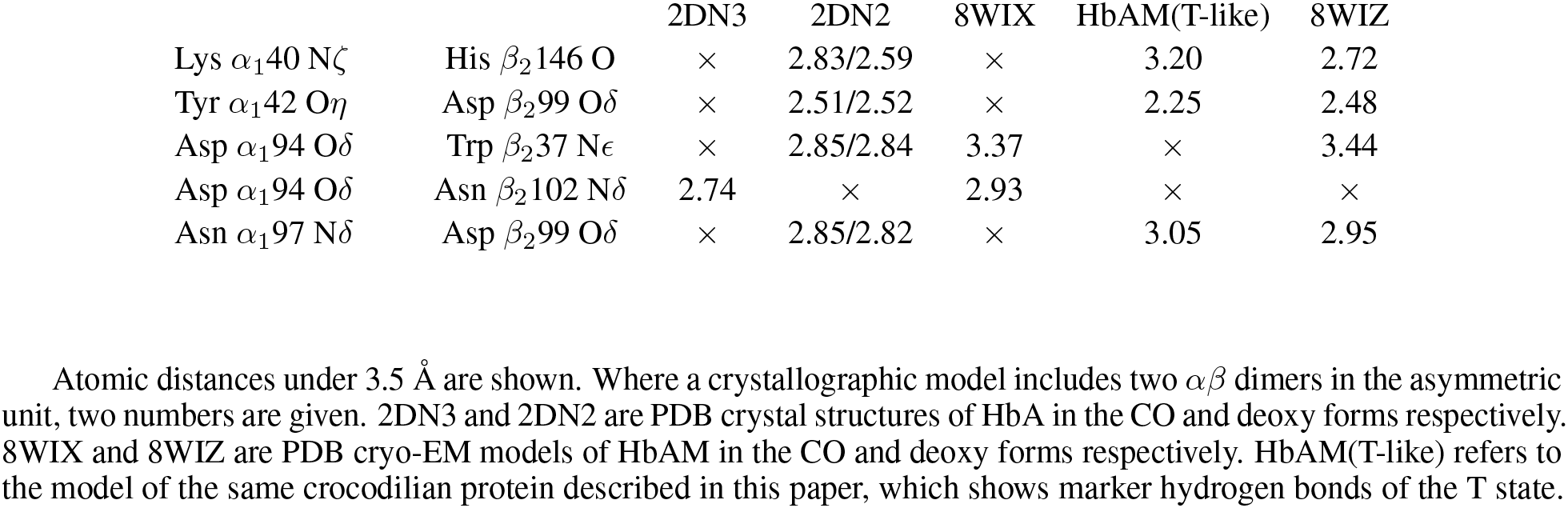
Various contacts between *αβ* dimers in different HbA and HbAM models.

